# Complex Spatial and Temporal Organization of Functional Brain Networks

**DOI:** 10.64898/2026.11.12.766207

**Authors:** Tyler A. Sassenberg, Sylia Wilson, Colin G. DeYoung

**Author notes:** Corresponding Author: Tyler A. Sassenberg, University of Minnesota Twin Cities, N218 Elliott Hall East River Parkway, Minneapolis, MN 55455.

## Abstract

Recent research exploring brain-behavior associations has relied on the language of complex systems to understand how stable behavioral individual differences emerge from brain function. At the same time, advances in neuroimaging have provided greater insight into both the topological and temporal properties of macroscale cortical networks. These new approaches offer a potential avenue for understanding how functional activity in the brain might behave in a manner that aligns with complex-systems interpretations of psychological traits. The present study used resting-state data from the Human Connectome project (*N* = 994) to explore associations between the spatial and temporal features of functional brain networks. Using an adaptive frequency approach to explore brain function at different timescales together with a family of methods from complex systems theory, we identified patterns of brain state organization reflecting a state-trait continuum, with more stable trait-like dynamics and greater functional coupling between networks in lower frequency oscillations. Additionally, we found evidence of greater coupling across timescales in brain networks associated with higher-order cognition. This research supports interpretations of brain organization as a spatiotemporal hierarchy and provides future avenues for exploring brain-behavior associations from a complex system perspective.

---

Research across psychology and neuroscience increasingly aims to explain how stable behavioral differences emerge from properties of the brain. A central challenge in this line of research is determining which neural features best map onto psychological traits. Recent theoretical work has drawn on principles of cybernetics (the study of goal-directed systems that self-regulate via feedback) and complex systems theory to explain how cognition and behavior emerge from patterns of brain function (DeYoung 2015; Friston et al. 2021; Safron and DeYoung 2021; Seth 2015). These theories outline how systems in the brain operate under specific conditions including goal-directed feedback, nonlinearity, and sensitivity to initial conditions (Hancock et al. 2022). These phenomena are hypothesized to emerge from the fundamental challenge of all biological organisms to manage metabolic costs and thermodynamic constraints under a framework known as active inference. Active inference describes the process through which embodied agents maintain persistent boundaries between themselves and their environment, either through updating generative models or acting in the environment (Friston et al. 2017). The accumulating evidence from active inference research indicates that brain function is organized to minimize prediction error, or the discrepancy between observed and expected states. From this perspective, identifying how error-minimizing processes are implemented in the brain in ways that differ systematically between people may help to explain patterns of behavioral individual differences, including cognitive abilities, normal range personality, and risk for psychopathology.

The concepts of cybernetics and complex systems have also been incorporated in theories of psychological individual differences. Psychological traits are commonly defined as density distributions of states, such that they reflect the probability of occupying particular states over time (Fleeson and Gallagher 2009). Explanatory accounts describe trait dimensions as reflecting between-person variation in parameters of a cybernetic system, or attractors in a dynamical landscape (DeYoung 2015; DeYoung and Krueger 2018; Sosnowska et al. 2019; Fajkowska 2015; Hovhannisyan and Vervaeke 2022; Hopwood et al. 2022; Olthof et al. 2023). From these perspectives, traits bias the system toward certain patterns of thought and behavior while still allowing variability in their expression. Because traits are probabilistic, rather than deterministic, individuals can deviate from these attractors in response to specific situational demands. Additionally, the attractor landscape itself can shift over time, such as during the onset of psychopathology, intervention, or normative development across the lifespan (DeYoung and Krueger 2018; Hudson and Fraley 2015; Bleidorn et al. 2018). An important implication is that patterns of behavioral individual differences span multiple timescales, where individuals experience fluctuations in psychological states at various durations, constrained by relatively stable dispositional traits (Andresen et al. 2024). Some patterns are purely state-like, others persist for more extended periods of time, and still others reflect patterns that are stable for many years or even decades.

This state-trait continuum presents an additional challenge to understanding the neural correlates of individual differences. Cybernetic theories of personality present hypotheses of how traits (including both normal-range personality differences and also persistent features of psychopathology) correspond to distinct cybernetic mechanisms encoded in functional brain networks and neurotransmitter systems (Allen and DeYoung 2017; DeYoung and Krueger 2018; DeYoung 2015), but it’s less clear how the properties of these neural mechanisms map onto the attractors used to describe traits, or how these traits change over time. Traits are moderately heritable, suggesting these attractors are grounded in neural architecture with a genetic basis (Schwaba et al. 2025), yet they can change through experience. Together, these points reflect the need to explore brain function across different timescales.

## Spatial and Temporal Aspects of Brain Function

Recent advances in neuroimaging provide greater understanding of both the spatial and temporal organization of brain function. Functional magnetic resonance imaging (fMRI) has been widely used to study large-scale functional brain organization due in part to its high spatial resolution compared to other neuroimaging methods. FMRI studies consistently identify macroscale brain networks based on correlated blood oxygen level dependent (BOLD) signals across the cortex (Smith et al. 2013; Yeo et al. 2011; Glasser et al. 2016; Schaefer et al. 2018; Du et al. 2024). These networks are organized hierarchically, with networks involved in self-related cognition and working memory being most functionally distinct from networks involved in sensory processing. This hierarchy is thought to arise from complex interactions of white matter tract projections, cortical folding, and localized neurotransmitter gradients (Atasoy et al. 2016; Buckner 2025; Glomb et al. 2021; Huntenburg et al. 2017; Margulies et al. 2016; Pang et al. 2023). Graph theoretical analyses further show that the brain is organized in terms of a small-world network, characterized by high clustering and high efficiency (Achard et al. 2006; Bassett and Bullmore 2006; Sporns and Zwi 2004; Bullmore and Sporns 2012). This small-world organization features hub nodes that adaptively consolidate information within functionally distinct modules, while simultaneously allowing the flexible exchange of information across modules (Bertolero et al. 2017; Crossley et al. 2014; van den Heuvel et al. 2011; Mengistu et al. 2016). Recent work has demonstrated that gradient and graph theoretical perspectives on the brain share core statistical features, reflecting a common pattern of brain organization (Rubinov 2025).

Temporal dynamics in brain activity can also be understood using principles from complex systems theory. In general, complex cybernetic systems exploit information bottlenecks like the hubs found in small-world networks to generate interpretable models of the environment (Goekoop and de Kleijn 2021; 2023, Hartl et al. 2026). In the active inference literature, this hierarchical hub configuration is described as an efficient mechanism for dissipating prediction error. Hierarchical processing implies higher-order hub nodes model derivations of the activity from downstream nodes. This integrative activity of hub nodes creates a vertical dimension in the network that encodes longer timescales and complex part-whole relationships. Ascending the hierarchy, these hubs are thought to model increasingly abstract goals or inferences that are used to quell lower-order prediction errors. From this perspective, the frequency decomposition from a brain network with a sophisticated generative model exhibits a preponderance of low-frequency oscillations from higher-order hubs that are effectively “explaining away” the prediction errors of lower-order nodes (Auksztulewicz et al. 2017; Alamia & VanRullen 2019; Michalareas et al. 2016). Attractors of low frequency dynamics constrain those of higher frequency dynamics, and this facilitates the emergence of temporal features that are characteristic of complex systems, including near-critical dynamics, metastability, and scale-invariance (Cocchi et al. 2017; Hengen and Shew 2024).

Task paradigms using electroencephalography (EEG) have supported this interpretation through the observation that lower frequencies of cortical travelling waves carry encodings of inferences, while higher frequencies carry information about prediction error (Bastos et al. 2020; Mayer et al. 2016). In the domain of fMRI, research has explored similar ideas through the role of dynamic functional activity and connectivity across different timescales. This line of research has shown that different networks coordinate according to signal dynamics at different timescales, with default, limbic, and frontoparietal networks assuming a higher-order integrating role (Bolton et al. 2020; Ito et al. 2020; Liégeois et al. 2019; Schaefer et al. 2014). Other researchers leveraging gradient and graph theoretical approaches also demonstrate associations between brain network organization and the temporal complexity found in a hierarchical configuration (Friston et al. 2021; Papo and Buldú 2024; Zamora-López et al. 2016).

## Current Research

Building on these ideas, we used resting-state fMRI to explore the associations between the spatial and temporal organization of functional brain networks to understand plausible neural mechanisms that could underlie stable psychological traits and their expression across different timescales. To accomplish this, we used complementary methods from dynamical systems theory to reconstruct properties of complex attractors derived from BOLD timeseries components across different cortical networks. We used these methods to test the general hypothesis that the complex behavior of these attractors would differ across networks and timescales, according to each network’s known functional role and the nature of different frequency domains in processing information. First, to extract information from the fMRI BOLD signal across timescales, we used an adaptive frequency method called multivariate variational mode decomposition (MVMD). This approach isolates noise from multiple biologically relevant signal components of the BOLD signal across different frequency domains. Next, to model these brain dynamics as complex systems, we used an approach called recurrence quantification analysis (RQA). RQA can be used to identify various properties of a dynamical system, including its predictability, stability, and the overall strength of the attractor. We then used similar models known as recurrence networks to assess the underlying topological organization of network attractors. Finally, to examine interactions across networks and timescales, we constructed multiplex recurrence networks, allowing us to assess how dynamics in one network or frequency band relate to those in another.

We explored three different kinds of associations: each brain network at a single timescale, each brain network across multiple timescales, and multiple brain networks at each timescale. Because low frequency oscillations are thought to encode trait-like phenomena in the active inference paradigm, we hypothesized that the attractor dynamics from lower frequency modes would exhibit more stable, predictable, and recurring brain states relative to the dynamics from attractors in state-like higher frequency modes. Additionally, considering multiplex recurrence networks of brain networks across modes, we hypothesized that greater cross-timescale coupling would be observed in the default, frontoparietal, and limbic networks, consistent with their integrative roles. Lastly, in multiplex recurrence networks comparing networks at a single timescale, we hypothesized that greater between-network coupling would be observed at lower frequencies, as multiple networks integrate information from different functional modalities to encode more abstract and temporally extended goals, and would decrease at higher frequencies, reflecting domain-specific processing.

## Methods

### Participants

A total of 994 participants (528 female) were selected from the 1200 young adult sample of the WU-Minn Consortium of the Human Connectome Project (HCP) (Van Essen et al. 2012). Initial HCP exclusion criteria included a history of severe psychiatric, neurological, or medical disorders. Participants were excluded from our subsequent analyses due to missing LR-encoded resting-state fMRI data from the first scanning session (*N* = 116) or poor-quality network boundaries following the individualized parcellation procedure (*N* = 90). Participants’ ages ranged from 22 to 37 years old (*M* = 28.7, *SD* = 3.7). All participants provided informed consent. Further information regarding the consent procedure is described by Van Essen et al. (2013). All study protocols were approved by the Institutional Review Board of Washington University in St. Louis.

### fMRI Data Acquisition and Preprocessing

fMRI data were acquired using a customized 3T Siemens Skyra scanner for all participants at Washington University in St. Louis. The present study used a single left-to-right phase encoded resting-state scan acquired using the following parameters: 72 axial slices; TR = 0.72 s; TE = 33 ms; flip angle = 52°; multiband acceleration factor = 8; voxel dimensions = 2 x 2 x 2 mm^3^; pixel bandwidth = 2,290 Hz. Additionally, high-resolution T1-weigthed MPRAGE structural images were acquired for anatomical surface registration with the following parameters: TR = 24 s; TE = 2.14 ms; flip angle = 8°; voxel dimensions = 0.7 x 0.7 x 0.7 mm^3^. Resting-state scans were preprocessed using the HCP minimal preprocessing pipeline and motion artifacts were removed using ICA-FIX (Burgess et al. 2016). Details of the HCP minimal preprocessing pipeline are described in greater depth in previous literature (Glasser et al. 2013; Ugurbil et al. 2013).

### Group Prior Individualized Parcellation

Functional network boundaries were identified using an individualized cortical parcellation approach. For each participant, ICA-denoised resting-state fMRI scans in subject-native space were first resampled to the fsaverage5 cortical surface mesh, and the BOLD timeseries at each vertex were normalized to zero mean and unit variance. The resulting participant surface data were then overlaid with a pre-defined group atlas with 400 functionally distinct regions (Schaefer et al. 2018) mapped to the 17-network atlas defined by Yeo et al. (2011). A Bayesian algorithm was applied to iteratively adjust parcel boundaries to maximize within-parcel homogeneity according to each participant’s unique patterns of functional connectivity (Chong et al. 2017). This algorithm was applied across 20 iterations so that participants had no more than one vertex on the cortical surface mesh changing its parcel identification on the final iteration, ensuring stable modifications of parcel boundaries. Through this process, each participant acquired a unique variant of a standard group-level atlas, such that the boundaries of the parcels of the initial atlas optimally reflected each individual’s patterns of functional network organization, while retaining their identity across participants.

### Multivariate Variational Mode Decomposition

To analyze patterns of functional activity across different timescales, we extracted oscillatory components using an adaptive frequency mode decomposition approach called multivariate variational mode decomposition (MVMD; Rehman and Aftab 2019). This data-driven approach assumes that a multivariate signal of interest, such as the BOLD timeseries from multiple brain networks, can be represented as a linear combination of amplitude- and frequency-modulated mode functions. MVMD aligns components of multivariate timeseries and extracts a set of oscillatory components corresponding to a data-derived central frequency across each channel. In other words, the mode functions acquired from the fMRI BOLD signal using MVMD represent distinct synchronized components that share a common central frequency across all brain networks. This approach accommodates individual differences in harmonic information without the specification of hard-coded parameters while also accounting for the non-linearity and non-stationarity inherent in fMRI data, and has been demonstrated to perform better than other bandpass filtering approaches (Rehman and Aftab 2019; Eriksen and Rehman 2023). In the present study, we averaged the timeseries of the regions within each of the 17 functional brain networks and applied MVMD to produce a set of mode functions for each network. In line with past work, we specified the extraction of 10 mode functions of the multivariate timeseries for each participant (Morante et al. 2025).

### Recurrence Quantification Analysis

The analysis pipeline using complex systems methods to test associations between functional brain networks and different timescales is illustrated in Figure 1. To assess the properties of each mode function as a complex system, we used an approach called recurrence quantification analysis (RQA). RQA is a method based on Takens’ embedding theorem (Takens 1980), which demonstrates that a higher-dimensional latent space can be reconstructed from a time-delayed representation of a single observed dimension of the system (e.g. a mode function derived from fMRI BOLD signal). The reconstruction of this latent space, also known as phase space, provides a spatial representation of the states visited by the system and captures the underlying dynamics that modulate the behavior of complex systems. The insight underlying RQA is that, because the behavior of any complex system is governed by the interactions of many unobserved variables, the dynamics of these unobserved variables can be inferred by identifying varieties of recurring patterns among successive lags of the single observed variable. Observing recurring or unique patterns in this phase space can provide information on the kinds of dynamical properties driving the original system. These patterns provide information about not only the value or intensity of a particular brain state, but also information about the sequence of states preceding that state. Numerical estimates of these properties can be calculated using a simplified two-dimensional representation called a recurrence plot. The recurrence plot visualizes recurring patterns within the system in matrix form, where points in diagonal or vertical lines represent distinct patterns of recurring states across time. Assessing sequences of isolated or clustered points provides information about the predictable or chaotic dynamics of a system. An example of the RQA embedding procedure is illustrated in Figure 2.

**Figure 1.**
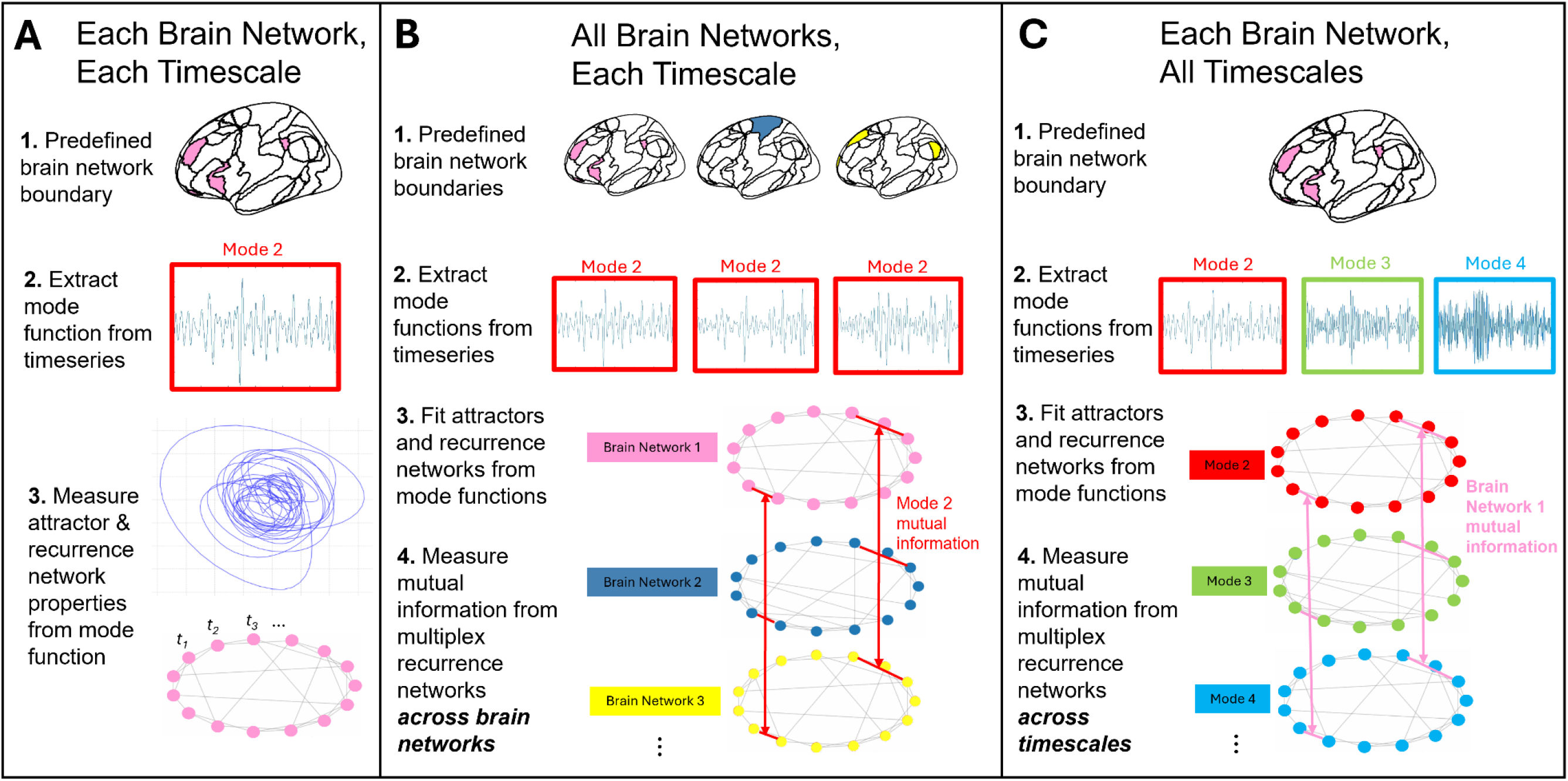
Methods used to test associations of complex dynamical properties of functional activity in each individual brain network at a given timescale (**A**), across all brain networks at a given timescale (**B**), and in each individual brain network across all timescales (**C**).

**Figure 2.**
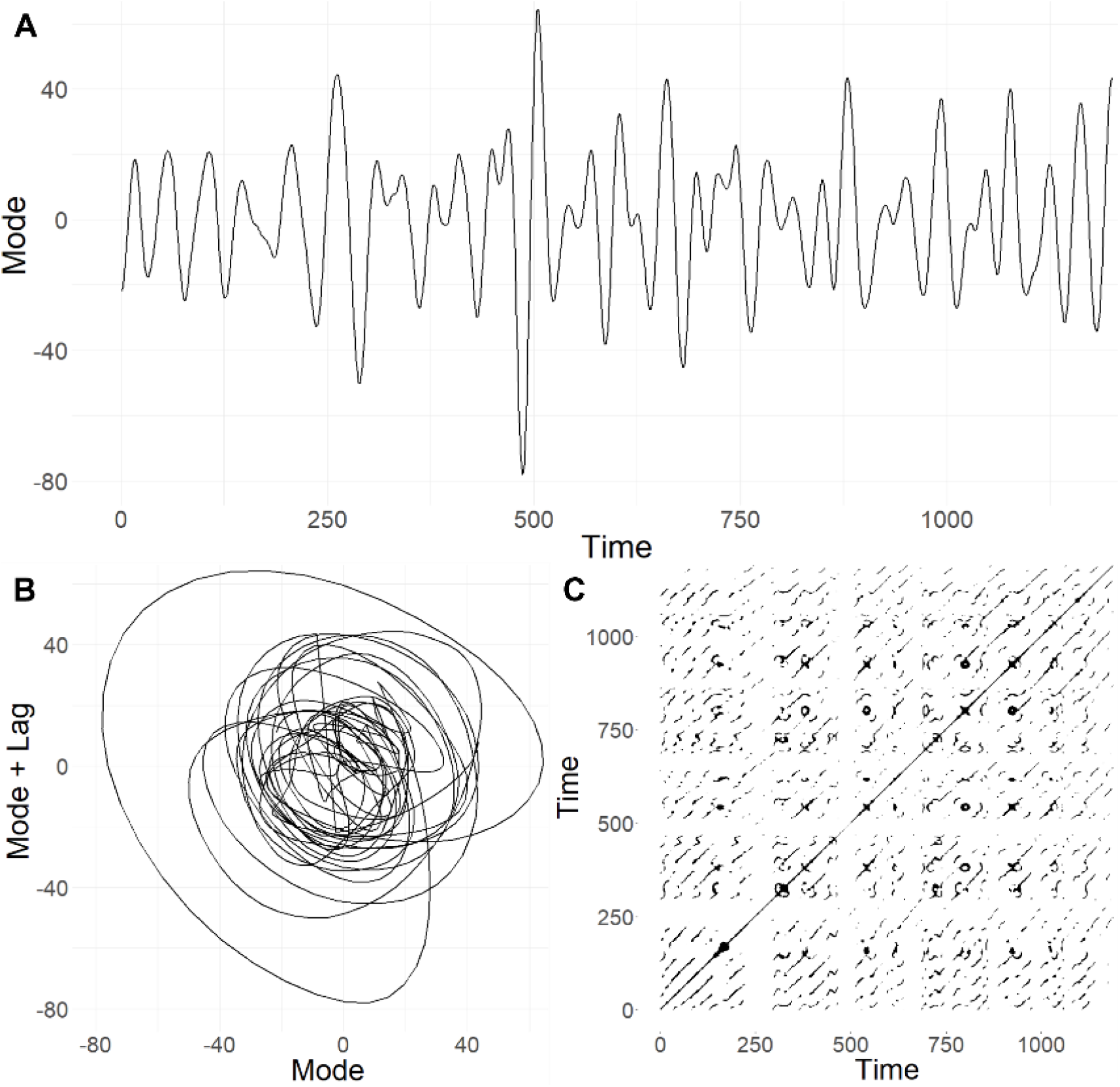
Recurrence quantification analysis procedure. (**A**) Mode function timeseries are derived using MVMD for each brain network across participants. (**B**) Delay embedding using false nearest neighbor analysis is used to reconstruct a higher-dimensional phase space from the mode functions. Trajectories through phase space represent the progression through various brain states over time. (**C**) Dynamical properties of these embedded trajectories can be quantified using a recurrence plot. Points in this plot represent recurring brain states in phase space. Structures in the plot aside from the main diagonal line, including smaller diagonal lines, whorls, or vertical lines, provide information about the dynamics of the brain states.

In the present study, we used RQA to fit a latent phase space for each functional network across each mode function for all participants. Phase space dynamics using RQA were calculated using the *casnet* package in R version 4.4.0 (Hasselman 2022; R Core Team 2024). With RQA, it is necessary to specify a lag, embedding dimension, and radius for the phase space embedding. These parameters are optimized empirically based on the timeseries data. To estimate the lag, the average mutual information across timeseries was calculated, and the first local minimum was selected for embedding. In this context, the first local minimum represents the number of time points after which a time-lagged version of the timeseries exhibits the greatest temporal independence from the original timeseries. Creating an embedding using this lag ensures that the time-lagged timeseries provide the most unique information (Wallot and Leonardi 2018). Next, the embedding dimension was selected using the first local minimum derived from false-nearest-neighbor analysis (Kennel et al. 1992). The embedding dimension refers to the number of surrogate dimensions needed to extract the dynamic properties of the underlying system. Lastly, in line with past research, we specified a fixed recurrence rate of 5% to ensure good comparability of phase spaces between individuals (Arellano-Véliz et al. 2024; van den Hoorn et al. 2020). Using RQA recurrence plots for each brain network across mode functions, we assessed six properties of complex dynamical systems: determinism, laminarity, mean diagonal line length, trapping time, diagonal line entropy, and vertical line entropy. These properties assess aspects of brain state dynamics in phase space, including the predictability, stability, length, and variability of different kinds of dynamics. Definitions of each of these metrics as well as their practical interpretations in the context of fMRI BOLD signal are described in Table 1. Detailed equations of these different metrics are described elsewhere (Marwan et al. 2007; Tommasini et al. 2022).

**Table 1.** Definitions and interpretations of recurrence quantification analysis metrics.

| Metric | Definition | Interpretation |
|---|---|---|
| Determinism | Proportion of recurring (≥2) points along *diagonal* lines in the recurrence plot, where line length or periods of recurrences vary. In phase space, deterministic patterns form adjacent, overlapping, or parallel trajectories. | Higher values indicate the presence of *recurring patterns* in brain dynamics. Lower values indicate irregular or unpredictable patterns. |
| Mean Diagonal Line Length | Mean of all *diagonal* lines in the recurrence plot. Estimate of the average time when two segments of a trajectory are close to each other. | Higher values indicate greater *stability of recurring patterns* of brain dynamics over time. Lower values indicate unstable or short-lived recurring patterns. |
| Diagonal Line Entropy | Shannon entropy of the distribution of *diagonal* line lengths in the recurrence plot. Quantifies the range of observed diagonal line lengths. | Higher values indicate greater *variability in the duration of recurring patterns* of brain dynamics. Lower values indicate brain state patterns of similar duration. |
| Laminarity | Proportion of recurring (≥2) points along *vertical* lines in the recurrence plot. Vertical lines indicate episodes in which the system occupies stable attractor states. | Higher values indicate the presence of *structured attractor states* with intermittent chaotic brain dynamics. Lower values indicate more chaotic brain dynamics overall. |
| Trapping Time | Analogous to Mean Diagonal Line Length, but the mean of all *vertical* lines in the recurrence plot. Estimate of the average time when the system occupies an attractor state. | Higher values indicate a *greater "pull" of stable dynamics*. Lower values indicate weaker attractor states. |
| Vertical Line Entropy | Analogous to Diagonal Line Entropy, but the Shannon entropy of the distribution of *vertical* line lengths in the recurrence plot. Quantifies the range of observed times in attractor states. | Higher values indicate greater *variability in the duration of trapping times*. Lower values indicate consistent periods of attractor pull. |

*Note*. Detailed equations of these different metrics are described elsewhere (Marwan et al. 2007; Tommasini et al. 2022).

### Recurrence Networks

In addition to assessing the dynamics of brain state trajectories through latent phase space, we also used recurrence networks to assess general geometric properties of the attractor landscape. A recurrence network is a graph theoretical interpretation of the information derived from the recurrence plot generated using RQA (Donner et al. 2010; Zou et al. 2019). Unlike a functional brain network measuring patterns of temporal synchrony across multiple brain regions, a recurrence network uses each ordered time point in phase space as a node in the graph, and connections between nodes as the recurrences identified using RQA. By specifying the connections between time points as edges in a network, graph theoretical measures can be used to measure general trends in recurrence patterns across different periods of time (Donges et al. 2012). While measures derived from the recurrence plot using RQA provide information about the dynamical properties of phase space, the recurrence network instead provides information about the overall geometric organization of the attractor landscape. In the present study we fit recurrence networks using the information derived from the phase space dynamics of each functional network and mode function for each participant. We assessed three properties of these recurrence networks: global efficiency, average local clustering, and network transitivity (Donner et al. 2010; Donner et al. 2011; Zou et al. 2019). Definitions of these metrics and their practical interpretations are described in Table 2.

**Table 2.**
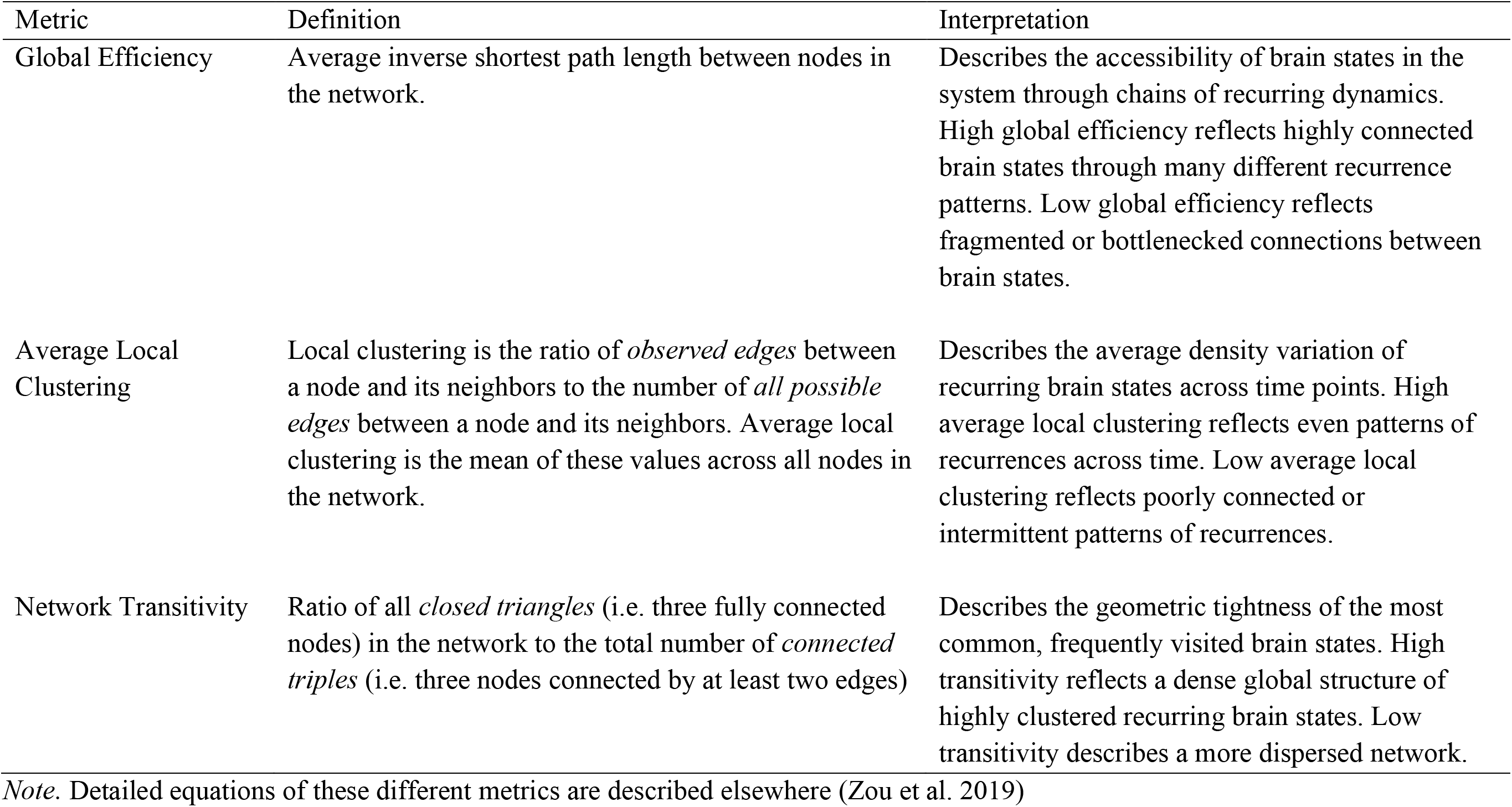
Definitions and interpretations of recurrence network metrics.

### Multiplex Recurrence Networks

Lastly, we used multiplex recurrence networks to assess patterns of communication between brain network attractors, as well as attractors across multiple timescales within a brain network. A multiplex network is a special case of a general multilayer network where each layer shares the same set of nodes but different types of edges can connect nodes both within and across layers (Boccaletti et al. 2014). A multiplex recurrence network is an extension of this idea, where each layer in the multilayer network is a recurrence network (Eroglu et al. 2018; Zou et al. 2019). Using these multiplex recurrence networks, we assessed the degree of communication across layers using a metric called inter-layer mutual information (IMI). IMI assesses the joint probability of the observed recurrences between identical sets of time points across the layers of the multiplex recurrence network. In other words, IMI indexes how much the structure of the attractor landscape is shared across different recurrence networks (Eroglu et al. 2018; Lacasa et al. 2015).

We specified two versions of these multiplex recurrence networks for each participant. The first version treated the recurrence networks of each functional brain network as separate layers to assess the shared dynamical properties between brain networks measured using a single mode function. For these multiplex recurrence networks, higher IMI indicates greater shared dynamical organization across functional brain networks at a given timescale. The second version used the recurrence networks of a single brain network assessed across each mode function as separate layers. For these multiplex recurrence networks, higher IMI indicates a greater degree of communication within a given brain network across multiple timescales. A necessary requirement of multiplex recurrence networks is that the recurrence networks across each layer contain the exact same nodes. To ensure this condition was satisfied for the multiplex recurrence networks comparing brain networks at a single timescale, we refit the phase space embeddings using the median embedding dimension for each brain network to reconstruct the recurrence networks in each layer. A similar approach was used for the multiplex recurrence networks comparing multiple timescales within a single brain network by using the median embedding dimension across each mode function.

### Analyses

We compared metrics derived from RQA, recurrence networks, and multiplex recurrence networks across functional brain networks and mode functions. First, to test the general properties of complex system metrics across timescales, we analyzed whole-cortex RQA and recurrence network metrics by averaging the values across the 17 networks for each mode function. We then tested differences in complex system metrics using a series of linear mixed-effects models specifying random intercepts for each participant. All models were fit using the *lme4*, and *lmerTest* packages in R, with denominator degrees of freedom adjusted using the Satterthwaite approximation (Bates et al. 2015; Kuznetsoca et al. 2017). We fit models predicting whole-cortex attractor dynamics for each metric across the four mode functions, as well as two-way models including main effects of mode function and brain network. Next, to test whether degrees of communication between brain network attractors differed across timescales, we fit another model to test mean differences in IMI from multiplex recurrence networks across mode functions. Lastly, to test whether brain network attractors differed in their communication across timescales, we fit a final model testing mean differences in IMI from multiplex recurrence networks across brain networks. All tests were corrected for Type I error using false discovery rate (Benjamini and Hochberg 1995).

## Results

### Mode Functions

Before assessing the dynamical features of each brain network, we assessed the central frequencies corresponding to each mode function, as well as their relative energy contribution, or the ratio of each mode’s energy to the overall energy of the signal. In line with past research using MVMD and similar approaches using fMRI data, we observed a dominant frequency (mode 1) centered around zero reflecting slow wave artifacts emerging from physiological noise, scanner instabilities, and head motion (Liu 2016; Power et al. 2011; Maknojia et al. 2019), as well as a set of mode functions (modes 2-5) within a neurophysiological range corresponding to frequencies attributable to the fMRI BOLD signal (Josephs and Henson 1999; Cordes et al. 2001). Modes 6 through 10 occupied a frequency range often attributed to fluctuations and harmonics of respiration, heartbeat, and cerebrospinal fluid pulsations (Frederick et al. 2012; Tong et al. 2019; Yuen et al. 2019). These mode functions are largely consistent with the frequency ranges previously reported in resting-state data from HCP (Morante et al. 2025). We also observed that the greatest proportion of the energy from the fMRI timeseries was derived from mode 2, representing the lowest end of the neurophysiological frequency range. We limited our subsequent analyses to mode functions 2-5 to assess functional properties specific to the BOLD signal across different frequency domains, while also removing non-BOLD oscillatory components found in modes 1 and 6-10. Distributions of the relative energy of each mode across participants are illustrated in Figure 3. The summary statistics of central frequencies for each mode are listed in Table 3.

**Figure 3.**
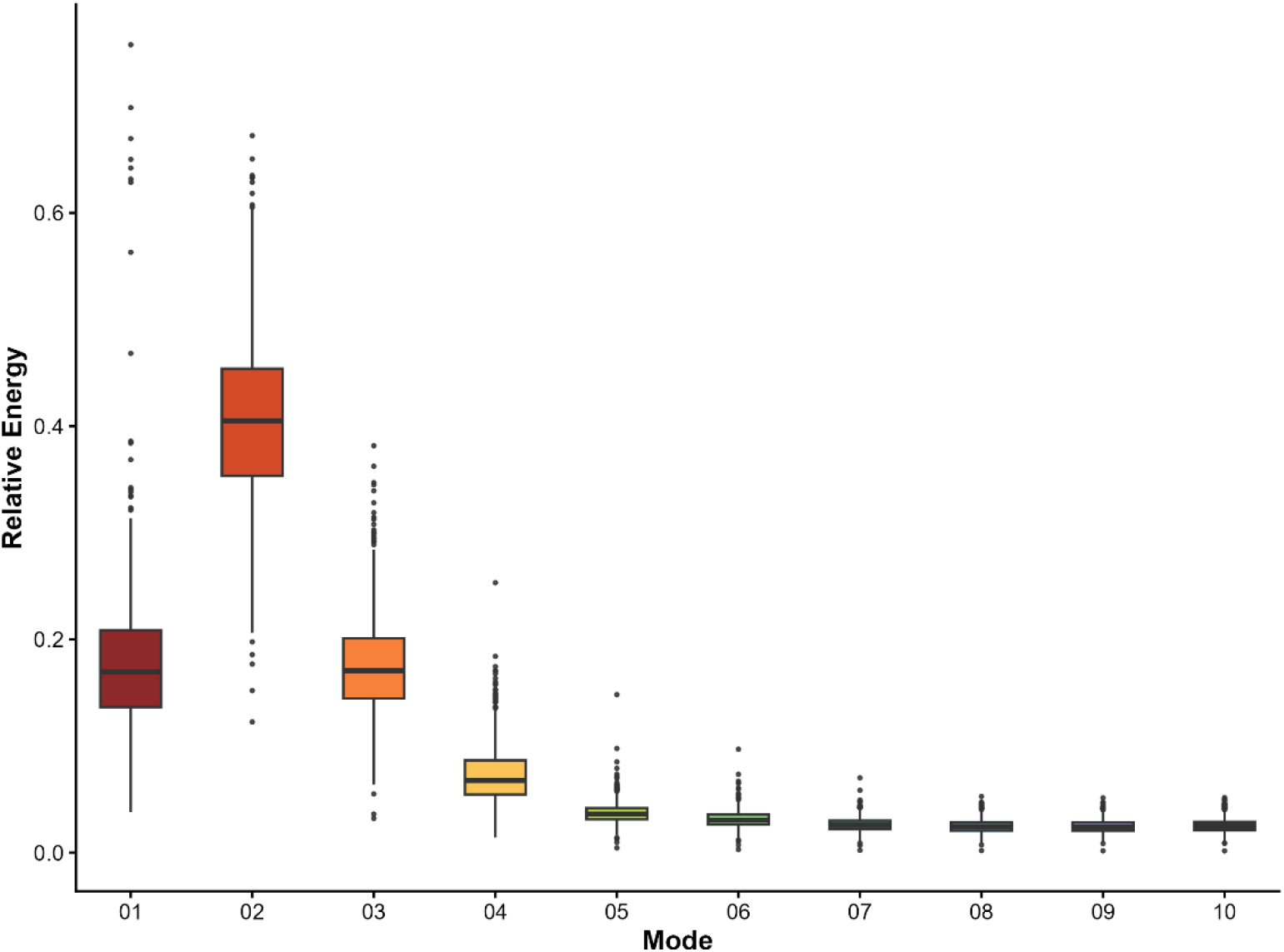
Whole-cortex relative energy distributions of each mode function across participants.

**Table 3.** Descriptive statistics of mode function central frequencies across participants.

| Central Frequency (mHz) | Mean (*SD*) | Skew | Kurtosis |
|---|---|---|---|
| Mode 1 | 0(0) | 0 | 0 |
| Mode 2 | 22.80(4.12) | .16 | -.12 |
| Mode 3 | 55.23(10.05) | -.19 | -.39 |
| Mode 4 | 99.00(15.49) | -.32 | -.36 |
| Mode 5 | 156.20(17.38) | -.73 | .41 |
| Mode 6 | 218.06(15.76) | -.84 | .72 |
| Mode 7 | 281.05(13.22) | -.93 | 1.04 |
| Mode 8 | 344.73(9.80) | -1.01 | 1.58 |
| Mode 9 | 408.58(7.75) | 2.42 | 23.77 |
| Mode 10 | 470.43(4.41) | -9.17 | 108.47 |

In addition to assessing the relative energy across the whole cortex, we compared the relative energy contributions of each mode function within each functional brain network. Relative energy profiles of each mode function across brain networks are illustrated in Figure 4. Consistent with the whole-cortex energy distributions, we found significant differences in the energy profiles across mode functions, where lower frequency modes exhibited higher energy compared to higher frequency modes (*F*[3,66579] = 97250.34, adjusted *p* < .0001). A significant main effect was found for individual brain networks (*F*[16,66579] = 344.33, adjusted *p* < .0001), where sensory networks exhibited a greater energy contribution from lower modes, while subnetworks of the default, frontoparietal, and limbic networks exhibited more even contributions across all mode functions. Individual main effects for each network and mode function in these models are reported in the supplemental material.

**Figure 4.**
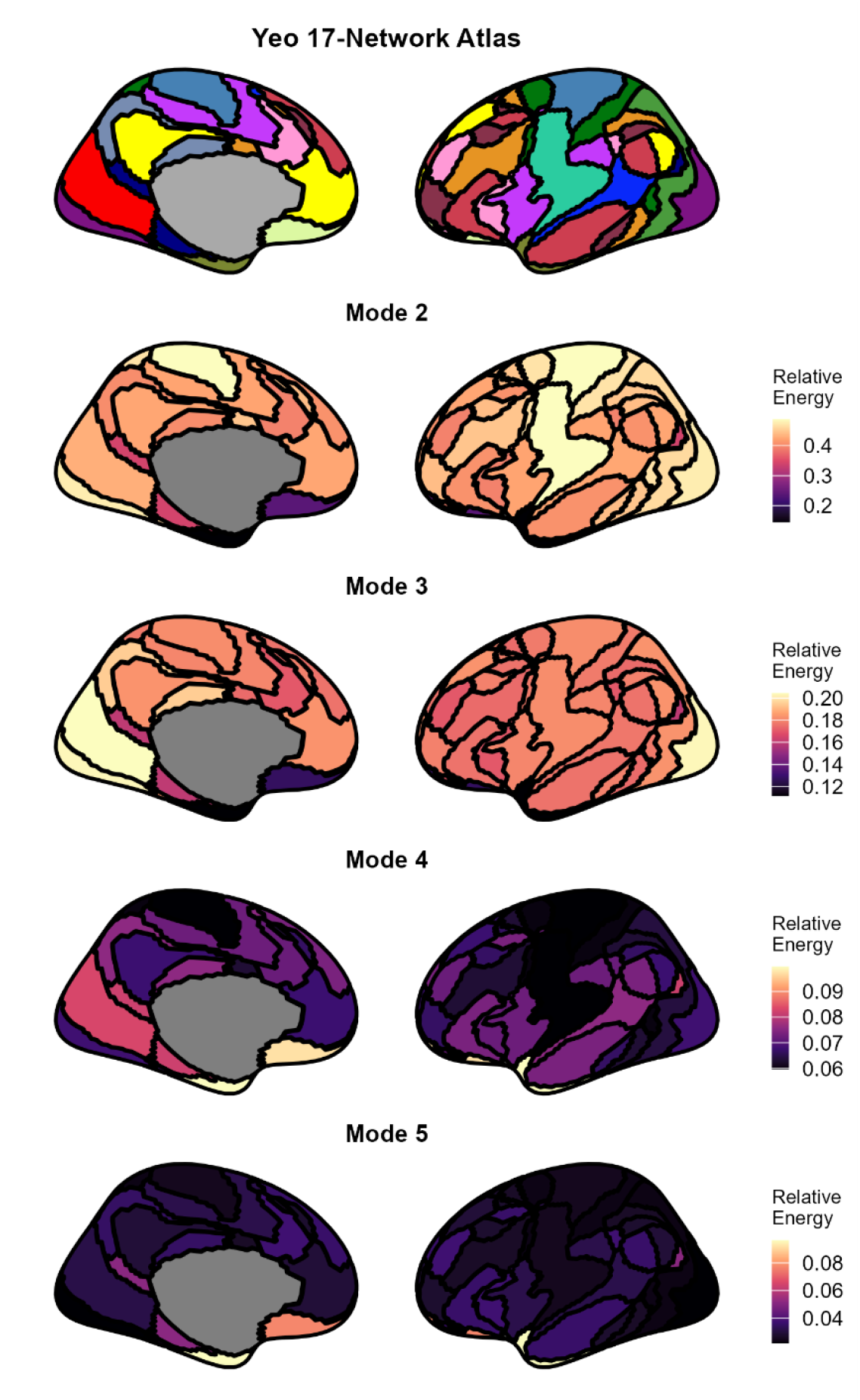
Mode function energy profiles in functional brain networks. Functional network boundaries were defined using individualized parcels corresponding to 17 networks defined by Yeo et al. (2011). Within each network, relative energy values sum to 1 across all 10 mode functions, but only mode functions 2-5 fall within frequency ranges corresponding to the BOLD signal.

**Table 4.** Omnibus F-tests of Mode Function Predicting Whole Cortex RQA Dynamics Metrics.

| Criterion | *F*-test | *p*FDR |
|---|---|---|
| Determinism | *F*(3,2979) = 7077.88 | <.0001 |
| Mean Diagonal Line Length | *F*(3,2979) = 2747.60 | <.0001 |
| Laminarity | *F*(3,2979) = 47519.88 | <.0001 |
| Trapping Time | *F*(3,2979) = 12462.55 | <.0001 |
| Diagonal Line Entropy | *F*(3,2979) = 2908.78 | <.0001 |
| Vertical Line Entropy | *F*(3,2979) = 34699.31 | <.0001 |

### Complex Brain Network Attractor Dynamics Across Timescales

Linear-mixed effects models revealed significant differences between each whole-cortex attractor dynamics metric across timescales. Omnibus *F*-test results of mode functions predicting each RQA metric are reported in Table 4, and main effects of these models are reported in the supplemental material. Boxplots of whole-cortex RQA metrics are illustrated in Figure 5. We observed significantly higher values for each of these measures in lower frequency mode functions, generally indicating attractors in lower frequency modes characterized by stable and highly predictable dynamics with recurring brain state sequences that are long but variable in length. Largely similar trends were observed in attractor dynamics of individual functional brain networks, with attractors of brain networks in lower frequency modes exhibiting greater signatures of stability and predictability, as well as small but significant differences in dynamical properties between networks. Omnibus *F*-tests of mode and brain network for each RQA metric are reported in Table 5. Individual main effects for each model are reported in the supplemental material.

**Figure 5.**
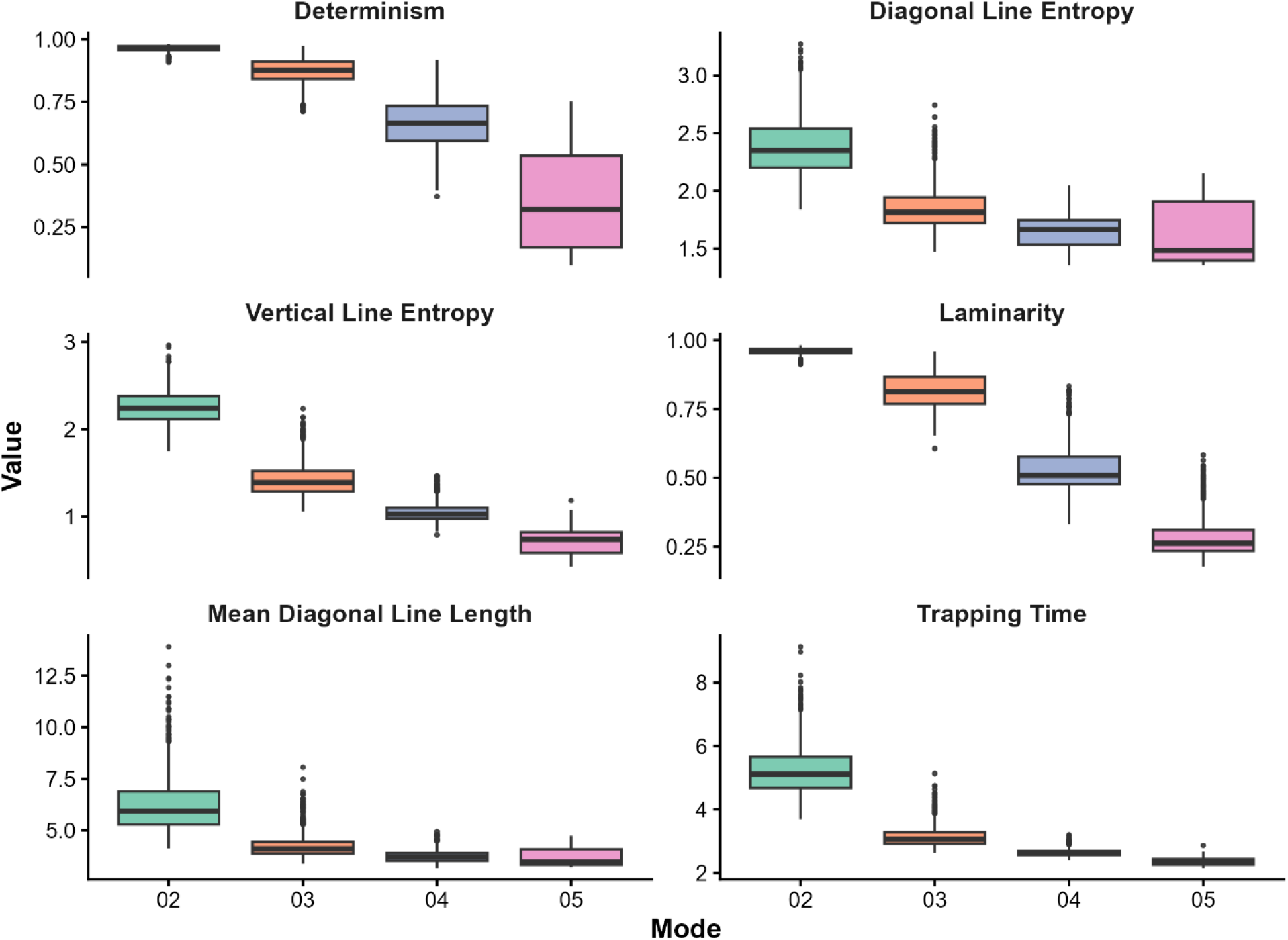
Distributions of whole-cortex attractor dynamics across mode functions for determinism, diagonal line entropy, vertical line entropy, laminarity, mean diagonal line length, and trapping time

**Table 5.**
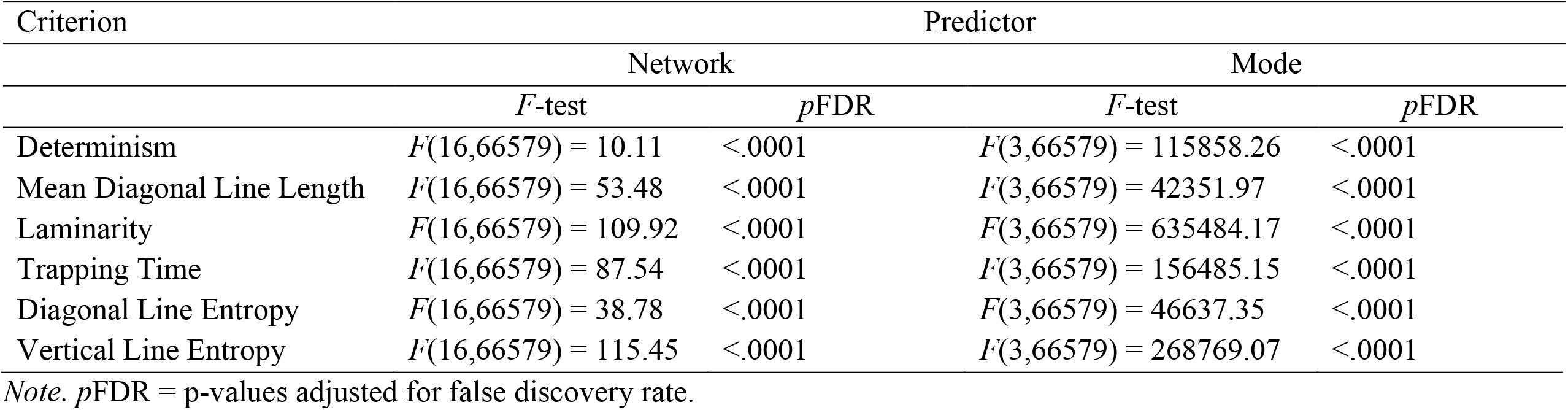
Omnibus F-tests of Brain Network and Mode Function Predicting RQA Dynamics Metrics.

### Complex Brain Network Attractor Geometry

In addition to the dynamics of brain states through phase space, we assessed the overall geometry of the attractor landscape using metrics derived from recurrence networks across the different mode functions. Omnibus *F*-tests of mode and brain network for attractor geometry metrics are reported in Table 6. Individual main effects for each model are reported in the supplemental material. Distributions of whole-cortex attractor geometry for each mode are illustrated in Figure 6. As with the RQA metrics assessing attractor dynamics, there were significant differences in patterns of whole-cortex global clustering (*F*[3,2979] = 410.34, adjusted *p* <.0001), global efficiency (*F*[3,2979] = 639.26, adjusted *p* <.0001) and network transitivity (*F*[3,2979] = 4817.97, adjusted *p* <.0001) across mode functions. Unlike the uniform patterns of attractor dynamics, we observed differences in features of attractor geometry across the mode functions, where higher global clustering was observed in both the highest and lowest frequency modes. Additionally, network transitivity was highest in the lowest frequency mode, and global efficiency was highest in the highest frequency mode. Similar patterns of attractor geometry were found for individual brain networks.

**Figure 6.**
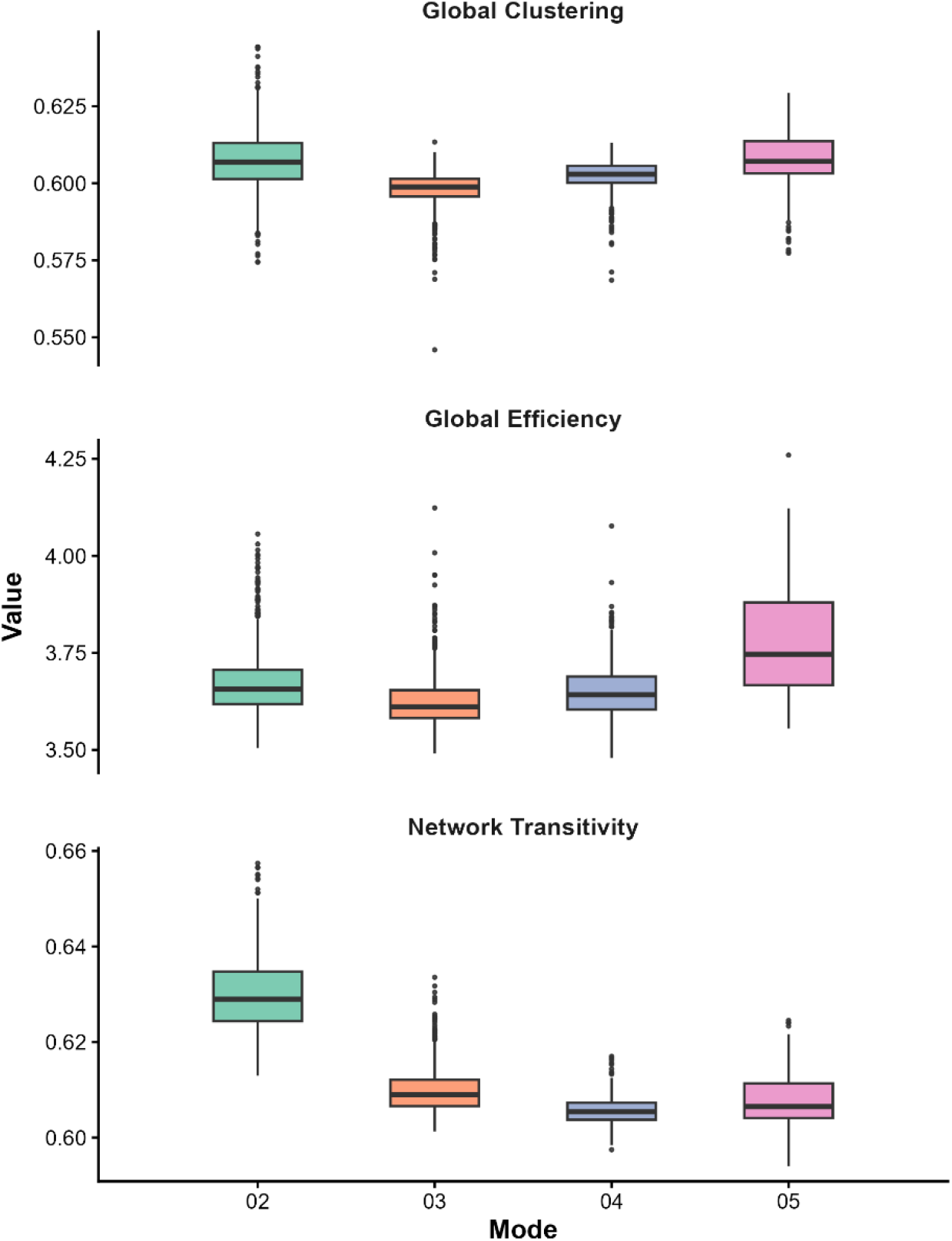
Distributions of whole-cortex global clustering (**A**), network transitivity (**B**), and global efficiency (**C**) derived from recurrence networks across mode functions.

**Table 6.** Omnibus F-tests of Brain Network and Mode Function Predicting Attractor Geometry Metrics.

| Criterion | Predictor |  |  |  |
|---|---|---|---|---|
|  | Network |  | Mode |  |
|  | *F*-test | *p*FDR | *F*-test | *p*FDR |
| Global Clustering | *F*(16,66579) = 104.69 | <.0001 | *F*(3,66579) = 2716.35 | <.0001 |
| Global Efficiency | *F*(16,66579) = 161.00 | <.0001 | *F*(3,66579) = 4952.56 | <.0001 |
| Network Transitivity | *F*(16,66579) = 41.13 | <.0001 | *F*(3,66579) = 18791.15 | <.0001 |

*Note*. *p*FDR = p-values adjusted for false discovery rate

### Inter-layer Mutual Information across Brain Networks and Timescales

Next, we analyzed the IMI from multiplex recurrence networks with layers of brain network attractors across each mode function. Distributions of IMI across mode functions and brain networks are illustrated in Figure 7. Individual effects for models of multiplex recurrence networks are reported in the supplemental material. A linear mixed effects model revealed a significant association of IMI across mode functions, with higher shared recurrences in the attractor landscape across networks in modes at lower frequencies (*F*[3,2979] = 57.61, adjusted *p* <.0001). Lastly, we tested differences in IMI from multiplex recurrence networks with layers of mode functions across each individual brain network. A linear mixed effects model revealed a significant association of IMI across functional brain networks (*F*[3,66582] = 345.16, adjusted *p* <.0001). We observed significantly higher levels of IMI within networks typically associated with higher-order cognition, including subnetworks of the default, frontoparietal, and limbic networks, with lower IMI across modes in unimodal visual and somatomotor networks.

**Figure 7.**
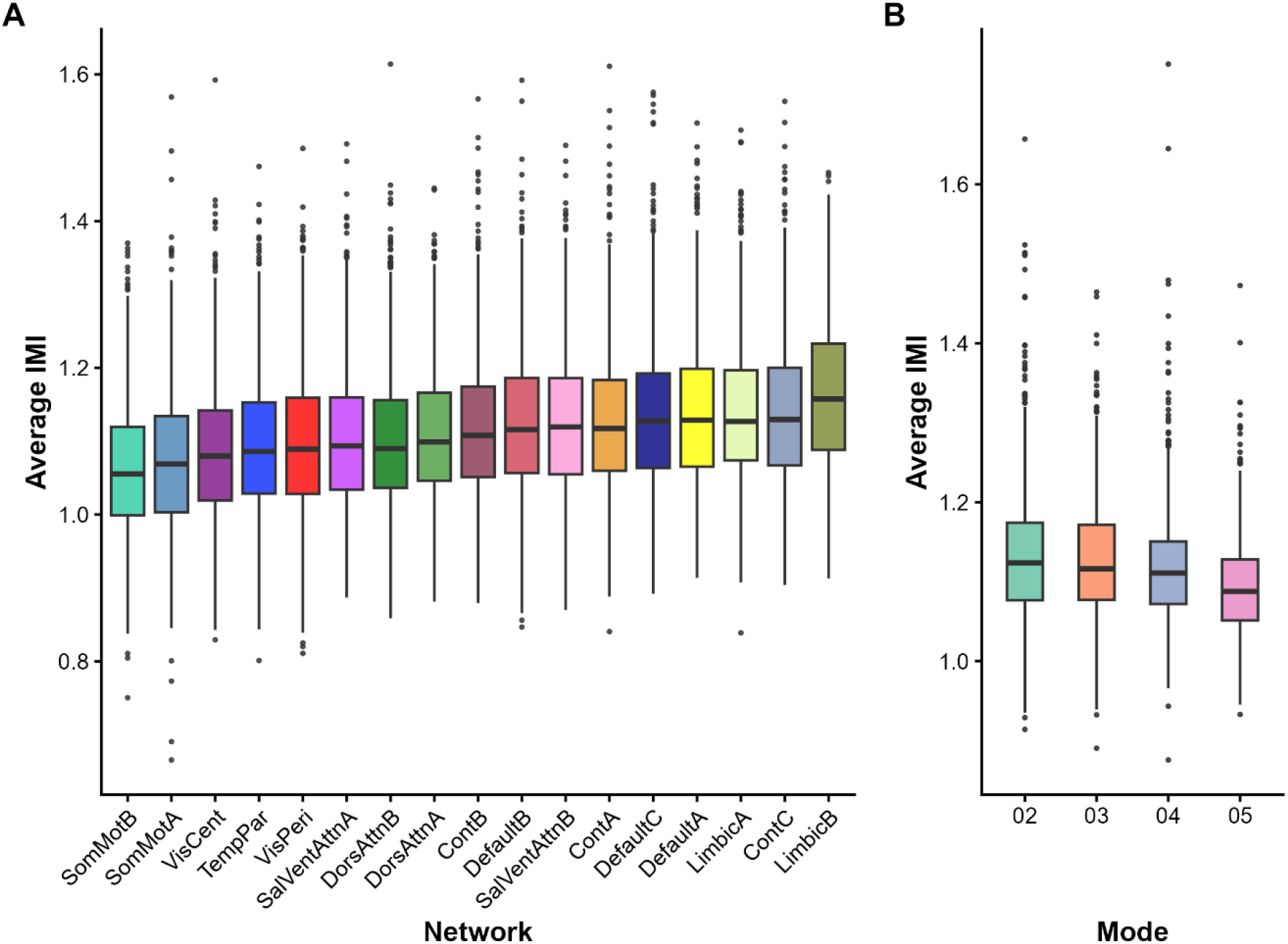
Distributions of average inter-layer mutual information (IMI) across multiplex recurrence networks. Average IMI between mode functions for each functional brain network are shown in (**A**). Average IMI between functional brain networks for each mode function are shown in **(B**). SomMot = somatomotor, VisCent = central visual, TempPar = temporoparietal, VisPeri = peripheral visual, SalVentAttn = salience/ventral attention, Cont = frontoparietal control.

## Discussion

Building on past research conceptualizing patterns of brain organization and individual differences in terms of complex systems, this research explored associations between the organization of functional brain networks across the cortex and across different timescales. Using resting-state fMRI in a large sample, we found evidence suggesting that the activity of functional brain networks exhibits features of complex systems in a manner that aligns conceptually with a state-trait continuum of brain activity. Specifically, these findings provide further evidence for different intrinsic timescales encoded in the brain’s functional activity, in line with past work assessing brain activity using dynamic functional connectivity and complex systems theory in fMRI data (Vidaurre et al. 2017; Vohryzek et al. 2020). We observed significant differences in complex features of dynamics across timescales. Lower frequency modes, hypothesized to reflect the encoding of trait-like goals within functional networks, exhibited higher values across a variety of complex system metrics. Together, these metrics provide a consistent profile of attractor behavior across timescales, where attractors in lower-frequency modes are more stable and predictable, indexed by higher determinism, laminarity, trapping time, and mean diagonal line length, but also exhibit greater variability in the length of recurring brain states, indicated by greater entropy. As the frequency increases, the attractors become less stable and more erratic, consistent with their hypothesized role in encoding situation-dependent information across short timescales.

These patterns of attractor dynamics are consistent with the different patterns of attractor geometry observed across different timescales. Rather than providing information about the momentary progression of observed brain states, the measures of global efficiency, global clustering, and network transitivity provide information about the underlying latent substrate guiding these dynamics. At the lowest frequency modes, attractors were characterized by a pattern of high clustering, high transitivity, and low efficiency. When organizing recurring brain states as a graph, this pattern indicates a well-connected set of brain states in a dense structure, but with certain sets of brain states being bottlenecked or rarely exhibiting transitions between dynamical regimes (Zou et al. 2019). In other words, this kind of attractor geometry steers brain states toward a highly clustered pattern of recurrences, with few accessible opportunities to deviate far from this pattern. This organization is reasonably consistent with the hypothesis that trait-like processing occurs at low frequency modes. In higher frequency modes, attractors tended to exhibit patterns of high clustering, low transitivity, and high efficiency. This pattern suggests a dispersed landscape of rapidly transitioning brain states, in contrast to the low frequency attractors.

In addition to the complex dynamics and organization of a single network, we also explored associations between networks at a given timescale. Using IMI as a metric of shared information in attractors between brain networks in multiplex recurrence networks, we found evidence of brain network attractors sharing more information at lower frequency modes compared to higher frequency modes. This observation supports our hypothesis that the encoding of more sophisticated, stable long-term goals and other inferences reflected in trait-like processing requires coordination across multiple information streams, whereas the information at higher frequency modes reflects more specialized processing. These findings are also consistent with work demonstrating that the emergence of a metastable regime that modulates chaotic dynamics occurs through the interactions of distinct functional modules in the brain (Moretti & Muñoz 2013; Palma-Espinosa et al. 2025). The greater degree of coupling in lower frequency domains of the BOLD signal is also consistent with past research suggesting that more typical functional connectivity metrics are primarily driven by, but not exclusive to, lower frequencies of the BOLD signal (Cordes et al. 2001; Niazy et al. 2011).

Additionally, when modeling different frequency modes as layers in the multiplex recurrence networks, we found that brain networks differ in their degree of coupling across timescales. Specifically, we found that attractors from subnetworks of the default, frontoparietal, and limbic networks exhibited greater shared information across timescales. This observation is consistent with past research reporting complex oscillatory patterns in these networks, suggesting that they occupy a fundamental role in integrating information across the brain (Deco et al. 2023; Krohn et al. 2023; Madan Mohan et al. 2026). At rest, this increased coupling could reasonably be viewed as an integrative function of reconciling short-term situational information with broader trait-like goals and inferences. This integration from a complex systems perspective also coheres with work showing how these networks exhibit unique profiles in processing information across timescales from an information theory perspective (Pope et al. 2025), as well as exhibiting distinct temporal metastates compared to sensory networks when assessed using dynamic functional connectivity (Vidaurre et al. 2017).

These findings provide a preliminary exploration of the spatial and temporal organization of functional brain networks from a complex systems perspective. The consistent patterns of attractor behavior aligns with past research describing functional activity organized along a state-trait continuum (Goekoop & de Kleijn 2021; 2023). The different patterns of complex features derived from timeseries across multiple scales have important implications for the nature of brain-behavior associations. Importantly, we did not test associations with behavioral individual differences in the present study. However, the organization we report in this study provides a wealth of potential avenues for linking functional brain activity to psychological traits according to theories describing these associations using the language of complex systems. The structure of attractors we report only provides information about brain states, but it is unclear whether the dynamics, underlying geometry, interactions of these attractors across networks or timescales, or some combination of these properties are most relevant for predicting psychological traits. We surmise that the stable features of attractors in low-frequency modes would be the strongest and most reliable predictors of behavioral individual differences. However, it may be just as important to compare individual differences in the more transient and chaotic dynamics of higher-frequency modes. Future research can explore these associations in greater detail, as well as pursue other aspects of complex organization, such as differences in complex organization between resting-state and task data, as well the predictive ability of these metrics in distinguishing normative from pathological trait variation, or their change through intervention or development.

### Strengths and Limitations

This work supports the use of a complex systems framework for understanding the complex spatial and temporal organization of the brain. We incorporated many forms of individualization in our analyses, including individualized brain network boundaries using GPIP, individualized harmonic information through the adaptive frequency procedure of MVMD, as well as the individually optimized phase spaces as part of RQA. These complementary approaches ensure precise measurement of both the spatial and temporal aspects of brain organization. Although past research has explored functional connectivity across timescales or complex organization of the overall fMRI signal (Kang et al. 2023; Krohn et al. 2023; Piccinini et al. 2021; Romanello et al. 2026), our work is unique in using these approaches together to assess a hierarchical configuration of complex attractor organization in the brain in a way that maps onto theoretical interpretations of broad trait dimensions.

However, there are several limitations to this study. First, although this work used a large sample with a relatively long scan, we incorporated information from only one resting-state run. To further expand on the nature of complex organization of the brain, future work would benefit from incorporating multiple runs of data to assess the reliability of each participant’s particular attractor landscape and dynamics, though past work has shown good reliability of complexity metrics in fMRI (Wehrheim et al. 2024). Additionally, functional networks differ in their temporal signal-to-noise ratio when assessed with common single-echo protocols (Ojemann et al. 1997; Girn et al. 2024), and it’s unclear to what extent that these differences are reflected in the mode functions derived from MVMD. Future research using multi-echo sequences can assess the extent to which this organization is reflecting unique scanning acquisition variability. Further, to limit the number of tests, we explored associations using only 17 functional networks as a single spatial scale. The framework we present can be expanded by incorporating the temporal decomposition of MVMD with complex systems methods across multiple spatial scales as well, including finer-grained analyses at the level of the individual voxel or functional parcel. Lastly, MVMD is a relatively new approach for analyzing the temporal features of fMRI data. Although it confers clear methodological advantages relative to other signal processing approaches, it is still unclear how the different mode functions in the neurophysiological range map onto distinct biological entities, and whether these entities are actual indirect reflections of neural activity or instead subtler features of vascular architecture (Tong et al. 2019; Yuen et al. 2019; Chen et al. 2020). For this reason, an approach like MVMD in fMRI might be enhanced in combination with more established temporal features of the brain like the canonical frequency bands found in EEG research.

## Conclusion

This research investigated associations within functional activity of brain networks from a complex systems perspective. Our results demonstrate that complex attractors derived from these brain networks exhibit significantly different properties across timescales, supporting theories of a state-trait continuum in brain organization. Attractors in lower-frequency modes exhibited more trait-like patterns of brain states compared to attractors in higher-frequency modes. Additionally, coupling between brain networks was observed in lower-frequency modes, and networks implicated in higher-order cognition exhibited greater integration across different timescales. These results are consistent with past interpretations of spatiotemporal brain organization, and provide a framework for testing more nuanced hypotheses about how patterns of behavioral individual differences relate to functionally distinct brain networks across different levels of temporal organization from the perspective of complex systems.

## Code/Data Availability

Resting-state fMRI data were obtained from the 2017 release of the HCP S1200. HCP has transitioned to the 2025 release as the primary distribution, but the legacy 2017 release is accessible through Amazon S3. Code used to produce individualized parcels of the Schaefer atlas using GPIP is available at https://neuroimageusc.github.io/GPIP. Code used to run MVMD is available at https://www.mathworks.com/matlabcentral/fileexchange/72814-multivariate-variational-mode-decomposition-mvmd. Code used in group level analyses is available at https://www.github.com/tylersassenberg/Brain-network-complexity.

## CRediT

**Tyler A. Sassenberg:** Conceptualization, Data curation, Formal Analysis, Investigation, Methodology, Project Administration, Software, Validation, Visualization, Writing – Original Draft, Writing – Review & Editing **Sylia Wilson:** Supervision, Methodology, Writing – Review & Editing **Colin G. DeYoung:** Methodology, Writing – Review & Editing

## Acknowledgements

Thank you to Maxwell Elliott for his comments on previous versions of the manuscript. Artificial intelligence (Claude Opus 5.5) was used to edit code used during data analysis and was not used to generate any text or images in this manuscript.

