## Supplemental Material for "Complex Spatial and Temporal Organization of Functional Brain Networks"

This supplementary material contains tables of coefficients of models describing differences in relative energy profiles across networks, as well as differences in RQA dynamics metrics, recurrence network properties, and multiplex recurrence network inter-layer mutual information (IMI).

### Table of Contents

Coefficients of models predicting relative energy from mode functions and brain networks 2

Coefficients of models predicting whole-cortex RQA dynamics metrics from mode functions 3

Coefficients of models predicting RQA dynamics metrics from mode functions and brain networks 4

Coefficients of models predicting whole-cortex recurrence network metrics from mode functions 7

Coefficients of models predicting recurrence network metrics from mode functions and brain networks 8

Coefficients of model predicting multiplex recurrence network inter-layer mutual information across timescales from brain networks 10

Coefficients of model predicting multiplex recurrence network inter-layer information across brain networks from mode functions 10

**Table A1**

*Coefficients of models predicting relative energy from mode functions and brain networks*

| Criterion | Predictor | *b*(SE) | *p*FDR |
|---|---|---|---|
| Relative Energy | Intercept (VisCent, Mode 2) | .425(.001) | <.0001 |
| | ContA | -.015(.002) | <.0001 |
| | ContB | -.014(.002) | <.0001 |
| | ContC | -.015(.002) | <.0001 |
| | DefaultA | -.016(.002) | <.0001 |
| | DefaultB | -.019(.002) | <.0001 |
| | DefaultC | -.034(.002) | <.0001 |
| | DorsAttnA | -.009(.002) | <.0001 |
| | DorsAttnB | -.010(.002) | <.0001 |
| | LimbicA | -.058(.002) | <.0001 |
| | LimbicB | -.079(.002) | <.0001 |
| | SalVentAttnA | -.015(.002) | <.0001 |
| | SalVentAttnB | -.024(.002) | <.0001 |
| | SomMotA | -.004(.002) | .019 |
| | SomMotB | -.004(.002) | .019 |
| | TempPar | -.015(.002) | <.0001 |
| | VisPeri | -.005(.002) | .002 |
| | Mode 3 | -.231(.001) | <.0001 |
| | Mode 4 | -.333(.001) | <.0001 |
| | Mode 5 | -.369(.001) | <.0001 |

**Table A2**

*Coefficients of models predicting whole-cortex RQA dynamics metrics from mode functions*

| Criterion | Predictor | *b*(SE) | *p*FDR |
|---|---|---|---|
| Determinism | Intercept (Mode 2) | .963(.003) | <.0001 |
| | Mode 3 | -.092(.005) | <.0001 |
| | Mode 4 | -.294(.005) | <.0001 |
| | Mode 5 | -.619(.005) | <.0001 |
| Mean Diagonal Line Length | Intercept (Mode 2) | 6.266(.025) | <.0001 |
| | Mode 3 | -2.032(.033) | <.0001 |
| | Mode 4 | -2.554(.033) | <.0001 |
| | Mode 5 | -2.624(.033) | <.0001 |
| Laminarity | Intercept (Mode 2) | .959(.002) | <.0001 |
| | Mode 3 | -.146(.002) | <.0001 |
| | Mode 4 | -.428(.002) | <.0001 |
| | Mode 5 | -.681(.002) | <.0001 |
| Diagonal Line Entropy | Intercept (Mode 2) | 2.383(.007) | <.0001 |
| | Mode 3 | -.535(.009) | <.0001 |
| | Mode 4 | -.734(.009) | <.0001 |
| | Mode 5 | -.771(.009) | <.0001 |
| Vertical Line Entropy | Intercept (Mode 2) | 2.252(.005) | <.0001 |
| | Mode 3 | -.828(.005) | <.0001 |
| | Mode 4 | -1.206(.005) | <.0001 |
| | Mode 5 | -1.539(.005) | <.0001 |
| Trapping Time | Intercept (Mode 2) | 5.234(.014) | <.0001 |
| | Mode 3 | -2.087(.017) | <.0001 |
| | Mode 4 | -2.597(.017) | <.0001 |
| | Mode 5 | -2.880(.017) | <.0001 |

**Table A3**

*Coefficients of models predicting RQA dynamics metrics from mode functions and brain networks*

| Criterion | Predictor | *b*(SE) | *p*FDR |
|---|---|---|---|
| Determinism | Intercept (VisCent, Mode 2) | .969(.003) | <.0001 |
| | ContA | -.005(.002) | .052 |
| | ContB | -.003(.002) | .202 |
| | ContC | -.007(.002) | .002 |
| | DefaultA | -.006(.002) | .010 |
| | DefaultB | -.005(.002) | .033 |
| | DefaultC | -.009(.002) | <.0001 |
| | DorsAttnA | -.001(.002) | .540 |
| | DorsAttnB | -.001(.002) | .572 |
| | LimbicA | -.015(.002) | <.0001 |
| | LimbicB | -.020(.002) | <.0001 |
| | SalVentAttnA | -.004(.002) | .138 |
| | SalVentAttnB | -.007(.002) | .006 |
| | SomMotA | -.001(.002) | .583 |
| | SomMotB | -.001(.002) | .681 |
| | TempPar | -.005(.002) | .059 |
| | VisPeri | -.004(.002) | .141 |
| | Mode 3 | -.092(.001) | <.0001 |
| | Mode 4 | -.294(.001) | <.0001 |
| | Mode 5 | -.619(.001) | <.0001 |
| Mean Diagonal Line Length | Intercept (VisCent, Mode 2) | 6.353(.02) | <.0001 |
| | ContA | -.086(.017) | <.0001 |
| | ContB | -.084(.017) | <.0001 |
| | ContC | -.132(.017) | <.0001 |
| | DefaultA | -.122(.017) | <.0001 |
| | DefaultB | -.085(.017) | <.0001 |
| | DefaultC | -.138(.017) | <.0001 |
| | DorsAttnA | -.048(.017) | .007 |
| | DorsAttnB | -.028(.017) | .119 |
| | LimbicA | -.220(.017) | <.0001 |
| | LimbicB | -.340(.017) | <.0001 |
| | SalVentAttnA | -.066(.017) | <.001 |
| | SalVentAttnB | -.103(.017) | <.0001 |
| | SomMotA | .027(.017) | .130 |

| | | | |
|---|---|---|---|
| | SomMotB | .024(.017) | .168 |
| | TempPar | -.059(.017) | <.001 |
| | VisPeri | -.031(.017) | .082 |
| | Mode 3 | -2.032(.008) | <.0001 |
| | Mode 4 | -2.554(.008) | <.0001 |
| | Mode 5 | -2.624(.008) | <.0001 |
| Laminarity | Intercept (VisCent, Mode 2) | .968(.002) | <.0001 |
| | ContA | -.008(.001) | <.0001 |
| | ContB | -.006(.001) | <.0001 |
| | ContC | -.014(.001) | <.0001 |
| | DefaultA | -.011(.001) | <.0001 |
| | DefaultB | -.010(.001) | <.0001 |
| | DefaultC | -.018(.001) | <.0001 |
| | DorsAttnA | -.004(.001) | <.001 |
| | DorsAttnB | -.002(.001) | .040 |
| | LimbicA | -.022(.001) | <.0001 |
| | LimbicB | -.028(.001) | <.0001 |
| | SalVentAttnA | -.006(.001) | <.0001 |
| | SalVentAttnB | -.011(.001) | <.0001 |
| | SomMotA | .002(.001) | .067 |
| | SomMotB | .003(.001) | .020 |
| | TempPar | -.006(.001) | <.0001 |
| | VisPeri | -.009(.001) | <.0001 |
| | Mode 3 | -.146(.001) | <.0001 |
| | Mode 4 | -.428(.001) | <.0001 |
| | Mode 5 | -.681(.001) | <.0001 |
| Diagonal Line Entropy | Intercept (VisCent, Mode 2) | 2.403(.005) | <.0001 |
| | ContA | -.018(.005) | <.0001 |
| | ContB | -.017(.005) | <.0001 |
| | ContC | -.028(.005) | <.0001 |
| | DefaultA | -.026(.005) | <.0001 |
| | DefaultB | -.019(.005) | <.0001 |
| | DefaultC | -.032(.005) | <.0001 |
| | DorsAttnA | -.009(.005) | .074 |
| | DorsAttnB | -.004(.005) | .401 |
| | LimbicA | -.052(.005) | <.0001 |
| | LimbicB | -.079(.005) | <.0001 |
| | SalVentAttnA | -.012(.005) | .012 |
| | SalVentAttnB | -.021(.005) | <.0001 |
| | SomMotA | .006(.005) | .202 |

| | | | |
|---|---|---|---|
| | SomMotB | .008(.005) | .106 |
| | TempPar | -.015(.005) | .003 |
| | VisPeri | -.010(.005) | .037 |
| | Mode 3 | -.535(.002) | <.0001 |
| | Mode 4 | -.734(.002) | <.0001 |
| | Mode 5 | -.771(.002) | <.0001 |
| Vertical Line Entropy | Intercept (VisCent, Mode 2) | 2.283(.005) | <.0001 |
| | ContA | -.038(.004) | <.0001 |
| | ContB | -.034(.004) | <.0001 |
| | ContC | -.057(.004) | <.0001 |
| | DefaultA | -.050(.004) | <.0001 |
| | DefaultB | -.037(.004) | <.0001 |
| | DefaultC | -.051(.004) | <.0001 |
| | DorsAttnA | -.020(.004) | <.0001 |
| | DorsAttnB | -.018(.004) | <.0001 |
| | LimbicA | -.062(.004) | <.0001 |
| | LimbicB | -.095(.004) | <.0001 |
| | SalVentAttnA | -.021(.004) | <.0001 |
| | SalVentAttnB | -.041(.004) | <.0001 |
| | SomMotA | .012(.004) | .002 |
| | SomMotB | .019(.004) | <.0001 |
| | TempPar | -.014(.004) | <.0001 |
| | VisPeri | -.020(.004) | <.0001 |
| | Mode 3 | -.828(.002) | <.0001 |
| | Mode 4 | -1.206(.002) | <.0001 |
| | Mode 5 | -1.539(.002) | <.0001 |
| Trapping Time | Intercept (VisCent, Mode 2) | 5.301(.012) | <.0001 |
| | ContA | -.084(.010) | <.0001 |
| | ContB | -.083(.010) | <.0001 |
| | ContC | -.131(.010) | <.0001 |
| | DefaultA | -.114(.010) | <.0001 |
| | DefaultB | -.077(.010) | <.0001 |
| | DefaultC | -.102(.010) | <.0001 |
| | DorsAttnA | -.044(.010) | <.0001 |
| | DorsAttnB | -.042(.010) | <.0001 |
| | LimbicA | -.130(.010) | <.0001 |
| | LimbicB | -.213(.010) | <.0001 |
| | SalVentAttnA | -.048(.010) | <.0001 |
| | SalVentAttnB | -.087(.010) | <.0001 |
| | SomMotA | .024(.010) | .013 |

| | | | |
|---|---|---|---|
| | SomMotB | .043(.010) | <.0001 |
| | TempPar | -.021(.010) | .034 |
| | VisPeri | -.024(.010) | .016 |
| | Mode 3 | -2.087(.005) | <.0001 |
| | Mode 4 | -2.597(.005) | <.0001 |
| | Mode 5 | -2.88(.005) | <.0001 |

**Table A4**

*Coefficients of models predicting whole-cortex recurrence network metrics from mode functions*

| Criterion | Predictor | *b*(SE) | *p*FDR |
|---|---|---|---|
| Global Clustering | Intercept (Mode 2) | .607(.001) | <.0001 |
| | Mode 3 | -.009(.001) | <.0001 |
| | Mode 4 | -.005(.001) | <.0001 |
| | Mode 5 | .001(.001) | .008 |
| Global Efficiency | Intercept (Mode 2) | 3.672(.003) | <.0001 |
| | Mode 3 | -.049(.004) | <.0001 |
| | Mode 4 | -.023(.004) | <.0001 |
| | Mode 5 | .103(.004) | <.0001 |
| Network Transitivity | Intercept (Mode 2) | .630(.001) | <.0001 |
| | Mode 3 | -.020(.001) | <.0001 |
| | Mode 4 | -.024(.001) | <.0001 |
| | Mode 5 | -.022(.001) | <.0001 |

**Table A5**

*Coefficients of models predicting recurrence network metrics from mode functions and brain networks*

| Criterion | Predictor | *b*(SE) | *p*FDR |
|---|---|---|---|
| Global Clustering | Intercept (VisCent, Mode 2) | .605(.001) | <.0001 |
| | ContA | .002(.001) | <.0001 |
| | ContB | .001(.001) | <.0001 |
| | ContC | .004(.001) | <.0001 |
| | DefaultA | .004(.001) | <.0001 |
| | DefaultB | .002(.001) | <.0001 |
| | DefaultC | .004(.001) | <.0001 |
| | DorsAttnA | .001(.001) | <.001 |
| | DorsAttnB | .001(.001) | <.001 |
| | LimbicA | .004(.001) | <.0001 |
| | LimbicB | .004(.001) | <.0001 |
| | SalVentAttnA | .002(.001) | <.0001 |
| | SalVentAttnB | .003(.001) | <.0001 |
| | SomMotA | -.001(.001) | .046 |
| | SomMotB | -.001(.001) | <.0001 |
| | TempPar | .001(.001) | <.0001 |
| | VisPeri | .002(.001) | <.0001 |
| | Mode 3 | -.009(.001) | <.0001 |
| | Mode 4 | -.005(.001) | <.0001 |
| | Mode 5 | .001(.001) | <.0001 |
| Global Efficiency | Intercept (VisCent, Mode 2) | 3.696(.003) | <.0001 |
| | ContA | -.031(.003) | <.0001 |
| | ContB | -.025(.003) | <.0001 |
| | ContC | -.054(.003) | <.0001 |
| | DefaultA | -.047(.003) | <.0001 |
| | DefaultB | -.026(.003) | <.0001 |
| | DefaultC | -.047(.003) | <.0001 |
| | DorsAttnA | -.018(.003) | <.0001 |
| | DorsAttnB | -.012(.003) | <.0001 |
| | LimbicA | -.050(.003) | <.0001 |
| | LimbicB | -.063(.003) | <.0001 |
| | SalVentAttnA | -.015(.003) | <.0001 |
| | SalVentAttnB | -.033(.003) | <.0001 |
| | SomMotA | .014(.003) | <.0001 |
| | SomMotB | .031(.003) | <.0001 |

| | | | |
|---|---|---|---|
| | TempPar | -.010(.003) | <.001 |
| | VisPeri | -.014(.003) | <.0001 |
| | Mode 3 | -.049(.001) | <.0001 |
| | Mode 4 | -.023(.001) | <.0001 |
| | Mode 5 | .103(.001) | <.0001 |
| Network Transitivity | Intercept (VisCent, Mode 2) | .631(.001) | <.0001 |
| | ContA | -.001(.001) | <.0001 |
| | ContB | -.001(.001) | <.0001 |
| | ContC | -.002(.001) | <.0001 |
| | DefaultA | -.002(.001) | <.0001 |
| | DefaultB | -.001(.001) | <.0001 |
| | DefaultC | -.002(.001) | <.0001 |
| | DorsAttnA | -.001(.001) | <.0001 |
| | DorsAttnB | .000(.001) | .093 |
| | LimbicA | -.002(.001) | <.0001 |
| | LimbicB | -.003(.001) | <.0001 |
| | SalVentAttnA | .000(.001) | .243 |
| | SalVentAttnB | -.001(.001) | <.0001 |
| | SomMotA | .001(.001) | <.0001 |
| | SomMotB | .001(.001) | <.0001 |
| | TempPar | .000(.001) | .603 |
| | VisPeri | .000(.001) | .179 |
| | Mode 3 | -.020(.001) | <.0001 |
| | Mode 4 | -.024(.001) | <.0001 |
| | Mode 5 | -.022(.001) | <.0001 |

**Table A6**

*Coefficients of model predicting multiplex recurrence network inter-layer mutual information across timescales from brain networks*

| Criterion | Predictor | *b*(SE) | *p*FDR |
|---|---|---|---|
| IMI | Intercept (VisCent) | 1.085(.003) | <.0001 |
| | ContA | .042(.004) | <.0001 |
| | ContB | .034(.004) | <.0001 |
| | ContC | .055(.004) | <.0001 |
| | DefaultA | .053(.004) | <.0001 |
| | DefaultB | .041(.004) | <.0001 |
| | DefaultC | .050(.004) | <.0001 |
| | DorsAttnA | .024(.004) | <.0001 |
| | DorsAttnB | .020(.004) | <.0001 |
| | LimbicA | .053(.004) | <.0001 |
| | LimbicB | .080(.004) | <.0001 |
| | SalVentAttnA | .018(.004) | <.0001 |
| | SalVentAttnB | .041(.004) | <.0001 |
| | SomMotA | -.012(.004) | .004 |
| | SomMotB | -.021(.004) | <.0001 |
| | TempPar | .010(.004) | .014 |
| | VisPeri | .012(.004) | .004 |

**Table A7**

*Coefficients of model predicting multiplex recurrence network inter-layer information across brain networks from mode functions*

| Criterion | Predictor | *b*(SE) | *p*FDR |
|---|---|---|---|
| IMI | Intercept (Mode 2) | 1.132(.002) | <.0001 |
| | Mode 3 | -.005(.003) | .092 |
| | Mode 4 | -.014(.003) | <.0001 |
| | Mode 5 | -.038(.003) | <.0001 |
